# Enhancing wheat yield potential through the *QFFE.perg-5A* and *QFEm.perg-3A* associated to spike fruiting efficiency: Insights from plot-level analysis

**DOI:** 10.1101/2024.08.14.607938

**Authors:** Nicole Pretini, Leonardo S. Vanzetti, Ignacio I. Terrile, Paula Silva, Giuliana Ferrari, Fernanda G. González

## Abstract

Wheat (*Triticum aestivum L.)* is a key global crop, essential for food security. To enhance its productivity this study focused on two quantitative trait loci (QTL), *QFFE.perg-5A* and *QFEm.perg-3A*, previously associated with fruiting efficiency at the spike level. We extended the analysis to the plot level to evaluate their impact on yield, thousand grain weight, and related traits, using a double haploid population of 102 lines in three environments. The results showed an epistatic interaction between a QTL for spike dry weight and grain number per spike. *QFFE.perg-5A* showed no interaction with the environment, increasing fruiting efficiency (13-15%), grain number per spike (7%), and spike number per unit area (9%), leading to an 8% rise in grain number per unit area and a 5% yield increase. In contrast, *QFEm.perg-3A*’s impact depended on the environment. This study connects spike-level QTL analysis with plot-level effects, providing valuable insights for breeding programs to improve wheat varieties with desirable traits and increased productivity.

## 1. Introduction

Wheat (*Triticum aestivum L.*) is one of the most important cereals supplying the global demand of food. Sustainable increases in its production is required to meet projected population growth (Borlaug, 2007; Chand, 2009; Reynolds et al., 2012; Fischer and Connor, 2018; FAO et al., 2021). Understanding the ecophysiological and genetic bases underlying potential yield can assist the breeding programs in enhancing crop productivity. However, yield is a complex trait determined by several components such as, grain number per spike (GNs), grain number per unit area (GN), thousand grain weight (GW_1000_), and spike number per unit area (SN) (Fig. 1). Analyzing numerical components in isolation often reveals trade-offs that can diminish or negate the benefits of enhancing individual traits. For instance, there are trade-offs between GNs and SN, where improvements in one trait may come at the expense of the other (Gaju et al., 2009; Dreccer et al., 2009, 2013). Therefore, it is necessary to consider an ecophysiological model that contemplates the relationships of growth and partition of assimilates among the different crop structures.

**Figure 1.**
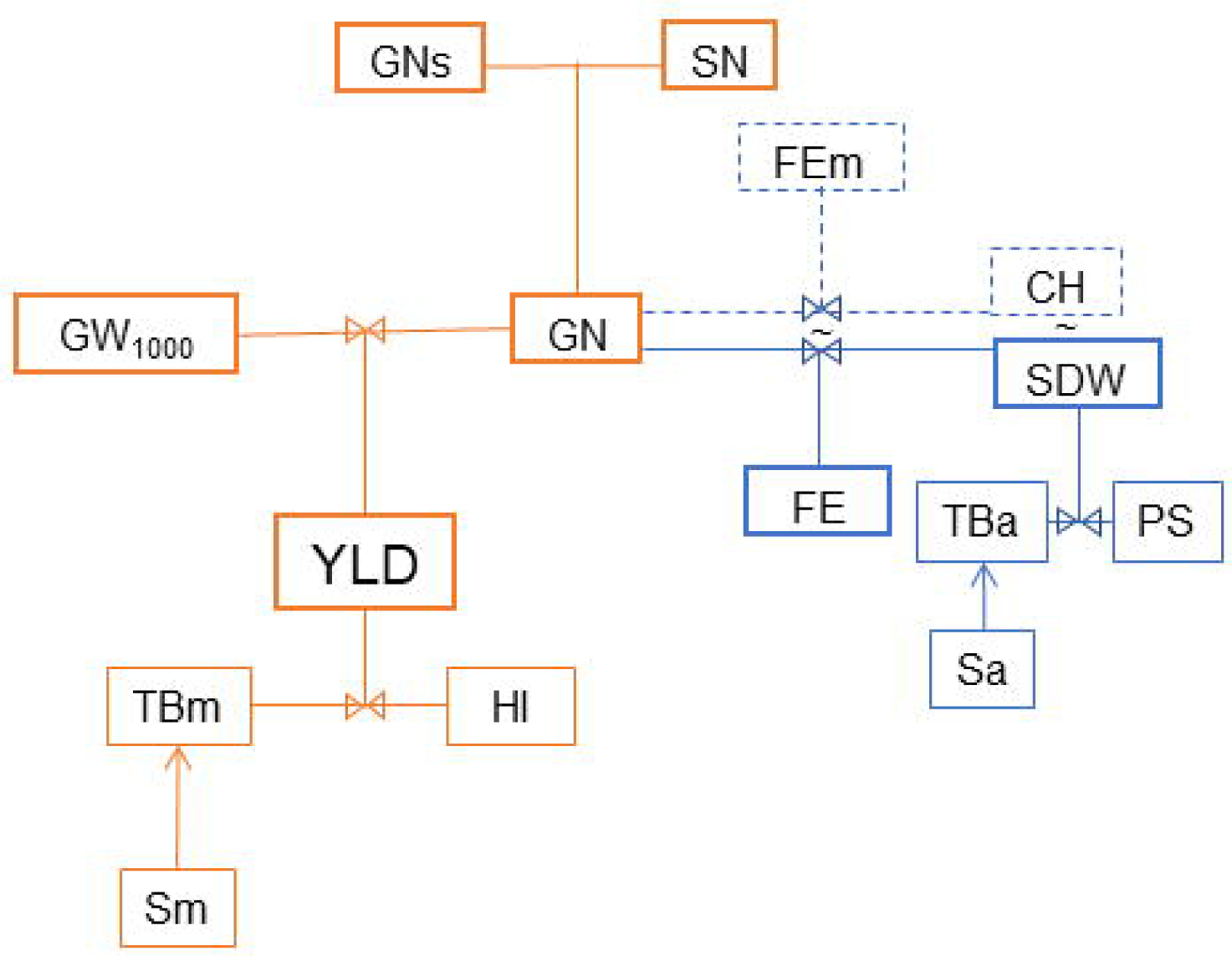
Main yield (YLD) components. Grain number per unit area (GN), thousand grain weight (GW_1000_), grain number per spike (GNs) and spike number per unit area (SN). The chaff (CH - nongrain spike dry weight at maturity) is generally used as surrogate for spike dry weight at anthesis (SDW) and the fruiting efficiency at maturity (FEm) is calculated instead of fruiting efficiency (FE). The relation of YLD with total biomass at maturity (TBm), stover (Sm) and harvest index (HI) is shown. Similarly the relation of SDW with total biomass at anthesis (TBa), partitioning of TBa to SDW (PS) and stover at anthesis (Sa) is presented.

The simplest way to understand yield considers the total biomass produced by the crop at maturity (TBm) and the portion of that biomass allocated to the reproductive organs, known as the harvest index (HI) (Fig. 1). This HI is associated with the amount of assimilates partitioned to the spikes around a critical period where yield is being determined. While yield formation occurs across the developmental stages, the timeframe spanning from 20 days before to 10 days after anthesis is critical for grain number determination (Fischer, 1975, 1985). The total biomass produced at anthesis (TBa) and the amount of assimilates partitioned to the spike during this period (PS) determines the final spike weight (SDW) and the fertile floret number at anthesis, and hence, the grain number at maturity (Fischer, 1975, 1985; Kirby, 1988; Ghiglione et al., 2008; González et al., 2011a).

Building upon this ecophysiological model, Fischer (1983) proposed an approach to understand the GN as the product of the spike dry weight at anthesis (SDW) and the fruiting efficiency (FE, the number of grains produced by unit of SDW - NG/SDW) (Fig. 1). FE is a promising trait for improving GN, and consequently enhancing yield, given the strong correlation observed between these variables (Abbate et al., 1998; González et al., 2011b). However, FE is a complex trait that integrates: (i) biomass partition within the spike between support structures and fertile florets, (ii) dynamics of floret evolution until the fertile floret stage, and (iii) grain set (grains per fertile floret). Consequently, FE can be conceptualized as the product between the fertile floret efficiency (FFE), representing fertile floret number (FF) established per unit of spike dry weight at anthesis (SDW), and grain set (GST, grains per floret) (Pretini et al., 2020a). Due to the complexity involved in directly measuring SDW, especially in large plot trials or early breeding generations, researchers often measure the dry weight of the spikes at maturity without grains (or chaff, CH) (Stapper and Fischer, 1990; González et al., 2011b; Martino et al., 2015; Alonso et al., 2018). Therefore, FE is commonly estimated as the grain number produced per chaff unit, referred as the fruiting efficiency at maturity (FEm), according to Fischer and Rebetzke (2018) (Fig. 1).

Molecular techniques have played a significant role in advancing plant breeding by enabling the identification and selection of traits related to defense and quality, offering a semi-quantitative approach. With the current abundance of available markers and the emergence of cost-effective, high-throughput platforms, there is a growing interest in leveraging these tools for more intricate and quantitative characteristics such as yield. The identification of new quantitative trait loci (QTL) associated with yield and its related traits serves as a valuable resource for breeding programs seeking to further enhance yield potential. To date, numerous studies have reported QTL mapping or fine mapping for yield and related traits (Quraishi et al., 2011; Chen et al., 2016, 2017; Deng et al., 2017; Cheng et al., 2017; Xu et al., 2017; Yu et al., 2018; Zhai et al., 2018; Pang et al., 2020; Isham et al., 2021). However, the translation of these QTL and associated markers into breeding programs has been limited, largely due to their minor effects and susceptibility to environmental influences, genetic background, and interactions (Cobb et al., 2019; Zheng et al., 2021). Also, most of these studies have identified numerical attributes of yield performance, with only a limited number addressing ecophysiological variables (Guo et al., 2017; Basile et al., 2019; Gerard et al., 2019; Pretini et al., 2021b). Furthermore, all these studies focused on detecting QTL for yield and associated traits at the individual spike level, without analyzing their impact at the plot-level.

In our previous study utilizing a DH population (Pretini et al., 2020b), we successfully identified and validated the presence of a QTL on chromosome 3A (*QFEm.perg-3A*) associated with fruiting efficiency at maturity at the spike level (FEms). Furthermore, we reported another QTL located on chromosome 5A (*QFFE.perg-5A*), which exhibited associations not only with the fruiting efficiency at spike level (FEs) but also with FEms and the fertile floret efficiency at spike level (FFEs). We thoroughly analyzed the effect of both QTL on the other traits associated to spike yield and found that the favorable allele of *QFEm.perg-3A* had no significant effect on FF or SDW per spike (SDWs), despite a slight increase in FFEs. However, *QFEm.perg-3A* did demonstrate a positive effect on FEs, resulting in greater grain number per spike (GNs), primarily explained by an increase in GST. Although, individual grain weight (GW) was reduced, yield efficiency (yield per spike/SDWs) increased, with a tendency toward higher yield per spike. Regarding *QFFE.perg-5A*, the presence of the favorable allele exhibited a positive effect on FFEs, leading to an increase in the number of FFs despite a reduction in SDWs. This allele also positively impacted GNs, attributed to increases in both FFs and GST. Despite a decrease in GW, there was an overall increase in yield efficiency, resulting in a higher yield per spike. To conclusively assess the utility of these QTL for breeding, it is essential to investigate their effects on yield and associated traits at the plot-level. This comprehensive approach will provide insights into potential trade-offs, such as those between GNs and spikes per m^2^ (SN) which are commonly observed.

The objective of this study was to investigate the influence of the quantitative trait loci *QFEm.perg*-3A and *QFFE.perg*-5A on the ecophysiological and numerical components determining wheat potential yield and associated traits, at the plot level.

## 2. Materials and methods

### 2.1. Plant materials and environments

The double haploid (DH) population utilized in this study was developed by González et al. (2011b). Comprising 102 lines, this DH population was created through the crossing of lines with contrasting FEm and FE, Baguette 19 with high and BioINTA 2002 with low (B19xB2002) according to González et al. (2011b) and Terrile et al. (2017).

The population was grown across three distinct environments, each characterized by a combination of location and year: (i) EEA Pergamino (33° 51′ S, 60° 56′ W) Research Station of INTA (Instituto Nacional de Tecnología Agropecuaria) during 2013 (P13) and 2015 (P15), and (ii) EEA Marcos Juárez (32° 72′ S, 62° 11′ W) Research Station of INTA during 2015 (MJ15). The experiments were conducted under no nutrient limitations (Phosphorus > 20 ppm, Nitrogen = 200 kg ha^-1^) and water constraints as rainfall was supplemented with irrigation from sowing to maturity. Precipitation levels varied across the different environments. The accumulated rain during the growth cycle was high in P15 (759 mm), while it was closer to the historical values in P13 and MJ15 (303 mm and 409 mm, respectively). In P13, irrigation amounted to 51 mm, reaching 354 mm from rainfall and irrigation. No irrigation was applied in P15 and MJ15. Weed, pest, and disease pressures were managed through appropriate chemical applications. Meteorological data was recorded at a station located less than 1000 m away from the experimental plots. For further details regarding experimental conditions, refer to Pretini et al. (2020a).

### 2.2. Experimental design and phenotyping

The population, along with both parents, was sown on the optimal date for each environment, following a randomized complete block design (RCBD) with two replications. Plots were arranged in five rows, each two meters long and spaced 0.21 meters apart. Plant density varied slightly across the different years: P13 had 330 plants per m^2^, while P15 and MJ15 had a slightly lower density of 280 plants per m^2^.

A half-meter section of the central row was sampled at anthesis (Z6.1, Zadoks et al., 1974) in P13 and P15, and spikes were separated from the rest of the biomass (or stover). The spike dry weight at anthesis (SDW, g m^-2^) and stover at anthesis (Sa, g m^-2^) were determined after drying in an oven at 70 °C for 72 h. The total biomass dry weight at anthesis (TBa, g m^-2^) was calculated as the sum of SDW and Sa. The partitioning of assimilates to the spike (PS) was estimated as the ratio of SDW to TBa

An additional half-meter section of a row was sample at maturity (Z9, Zadoks et al., 1974) in all three environments. Spikes were separated from stover, and their count was used to calculate the spikes per m^2^ (SN). After drying at 70 °C for 72 h, spikes were weighted and manually threshed to separate the chaff from the grains. The stover was also weighted to estimate its value per m^2^, Sm (g m^-2^). Grains were weighed before counting to estimate the grain yield (YLD, g m^-2^). Two subsamples of 2 grams each were counted in an automatic seed counter (Pfeuffer GmbH, 15-230 V, 50/60) to determine the thousand grain weight (GW_1000_). The grain number (GN, number m^-2^) was calculated as the ratio of YLD to GW_1000_. Chaff (CH, g m^-2^) was estimated by subtracting the weight of grains from the spike weight at maturity. The total of biomass at maturity (TBm, g m^-2^) was estimated as the sum of CH, Sm, and YLD. The harvest index was calculated as the ratio of YLD to TBm. The FE was estimated as the ratio of GN to SDW, while the FEm was calculated as the ratio of GN to CH. Finally, the grain number per spike (GNs) was computed as the ratio of GN to SN. The data at spike level used for these calculations was taken from Pretini et al. (2020a)

### 2.3. Data Analysis

The phenotype performance of the DH population and both parents was calculated across the two replicates in each environment. Descriptive statistics including population means, minimums, maximums, average, and standard deviation were estimated using the software InfoStat/P (Di Rienzo et al., 2010) to provide an overview of population distributions. A modified version of the Shapiro-Wilks test (Mahibbur and Govindarajulu, 1997) and a quantile-quantile (Q-Q) plot using residual values were employed to assess the normal distribution of the data. Person’s Correlation coefficients were calculated within and across environments to determine statistical relationships between variables and to explore potential trade-offs among them.

To assess the impact of *QFEm.perg-3A* and *QFFE.perg-5A*, on the 13 traits (SDW, Sa, TBa, CH, Sm, TBm, SN, GN, GW_1000_, YLD, HI, FEm, and FE) a factorial analysis of variance (ANOVA) was performed. The SNP peak marker for each QTL (*wsnp_CAP11_rep_c4226_1995152* for *QFEm.perg-3A* and *BS00083507_51* for *QFFE.perg-5A*) was used as class variables in the model. Additionally, two-way interactions were included to determine significant epistatic interactions between both QTL and the environment.

Genetic components of variance were estimated for YLD, GN, GW_1000_, GNs, SN, SDW, CH, FEm, and FE through ANOVA analyses, considering each environment as the combination of year and location, with blocks nested within the environments.

## 3. Results

### 3.1. Phenotypic performance of the DH population across environments

The mean temperatures during the reproductive period, from stem elongation to anthesis (September to October), varied across environments: ranging from 13.2 to 17.6 °C in P13, from 13.6 to 14.6 °C in P15, and from 14.9 to 20.8 °C in MJ15. Anthesis dates, within ± 4 days for each environment, were October 26 for P13, October 20 for P15, and October 11 for MJ15. The average temperature during grain filling in November was 20.3 °C in P13, 19.2 °C in P15, and 20.9 °C in MJ15. Further details can be found in Pretini et al., 2020a. All variables exhibited a normal or near-normal distribution within each specific environment, and transgressive segregation from the parental lines was observed (Fig. 2, Supplementary Table 1).

**Figure 2.**
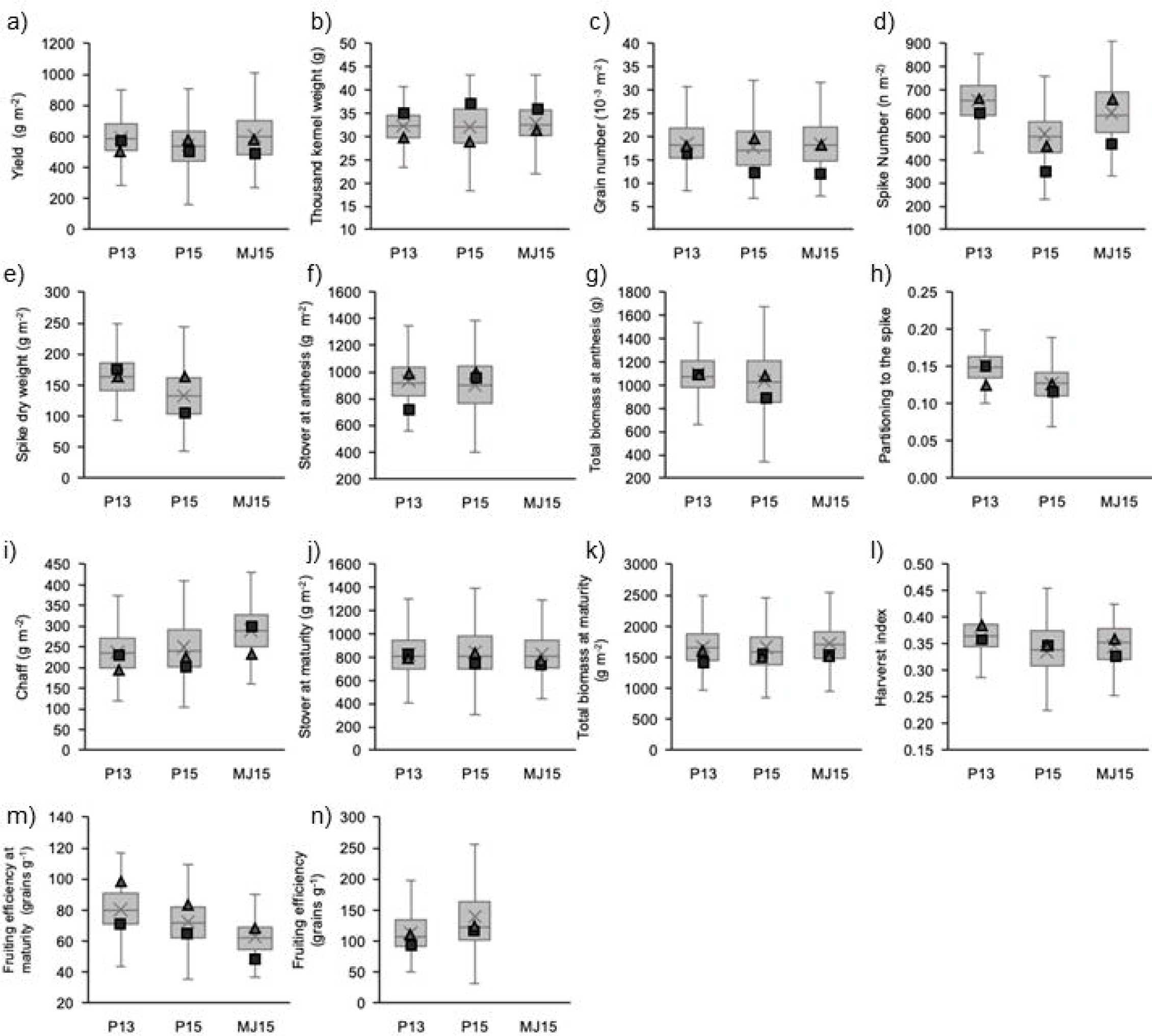
Boxplots of the traits studied in the DH population. B19 (triangles) and B2002 (squares) values are indicated in each plot. The bottom end represents the first/lower quartile, the line inside the box represents the median, the top end represents the third/upper quartile of the data. The cross represents the mean of the data. a) YLD: yield (g m^-2^), b) GW_1000_: thousand grain weight (g), c) GN: grain number (grains m^-2^), d) SN: spike number (spikes m^-2^), e) SDW: spike dry weight (g m^-2^), f) Sa: stover at anthesis (g m^-2^), g) TBa: total biomass weight at anthesis (g m^-2^), h) PS: partitioning to the spike, i) CH: chaff (no-grain spike dry weight at maturity, g m^-2^), j) Sm: stover at maturity (g m^-2^), k) TBm: total biomass weight at maturity (g m^-2^), l) HI: harvest index, m) FEm: fruiting efficiency at maturity (grains g_CH_^-1^), n) FE: fruiting efficiency (grains g_SDW_^−1^).

The mean YLD of the DH population showed 9% variation among environments, ranging from 555 to 603 g m^-2^ (Fig. 2a). Meanwhile the mean GN and GW_1000_ showed 6 and 3% variation (17551 to 18530 grains m^-2^ and 32.1 to 32.9 g, Fig. 2b and 2c). In contrast, the mean GNs observations from Pretini et al. (2020a) indicated higher GNs produced in P15 (56.4 grains spike^-1^), followed by MJ15 (42.8 grains spike^-1^) and P13 (36.4 grains spike^-1^). The mean SN was behind those differences, being lower in P15 (511 spikes m^-2^), intermediate in MJ15 (603 spikes m^-2^) and higher in P13 (654 spikes m^-2^, Fig. 2d). The SDW mean values showed a maximum of 20% variation (133 to 165 g m^-2^ Fig. 2e) while Sa, TBa and PS varied 15 to 13% between extreme environments (Fig. 2f, 2g, 2h). Similarly to the SDW, the mean CH changed 18% considering the different environments (Fig. 2i), and it was heavier than the SDW. The Sm, TBm and HI varied 3, 4 and 6% (Fig. 2j, 2k, 2l). Regarding the fertility indexes, the mean FEm varied from 63 grains g_CH_^−1^ to 80 grains g_CH_ ^−1^, while the mean FE ranged from 114 grains g_SDW_ ^−1^ to 140 grains g_SDW_ ^−1^ (Fig. 2l, 2m).

### 3.2. Relationship between yield and main components

According to the ANOVA results, YLD showed significant differences among genotypes and a significant genotype x environment interaction, with a relatively minor influence from the environment itself (Table 1). Genotypic effects accounted for 26% of the variation in YLD, while genotype x environment interaction explained 37% of the variation (Table 1). Similarly, the GN exhibited a minor effect from the environment (1% variation explained), but a significant effect from genotype and genotype x environment interaction, accounting for 29% and 38% of the variation, respectively (Table 1). For GW_1000_, once again, the environment had a small effect but it was highly significant, explaining 1% of the variation, while the genotype and genotype x environment interaction explained 63% and 21% of the variation, respectively (Table 1).

**Table 1.**
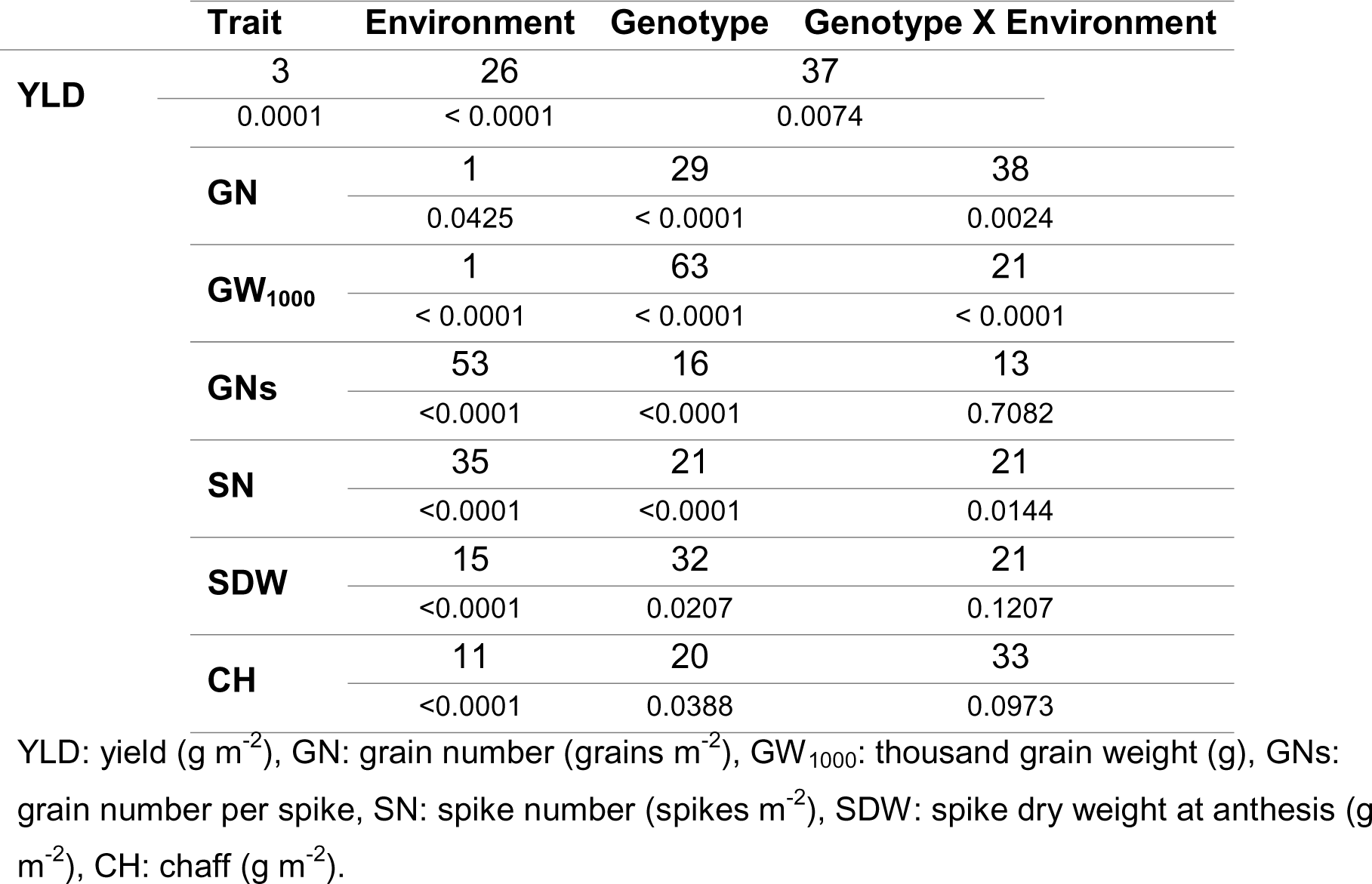
Percentage of total variation (considering the sum of squares) and the p-values for the environment, genotype, and genotype x environment interaction.

The variation in GNs was primarily explained by the environment (53%) and the genotype (16%), with no significant interaction between them. For the SN, the environment accounted for 35% of the observed variation, while both genotype and the genotype x environment interaction explained 21% (Table 1). The SDW variation was predominantly explained by genotype (32%), followed by the environment (15%) with no significant interaction. Similarly, for CH, no significant interaction was observed, with genotype explaining 20% and environment 11% of the variation (Table 1).

The phenotypic association between YLD and GN and GW_1000_ across the three environments was analyzed. When considering the entire subset of data, a strong positive correlation was observed between YLD and GN within and across environments (Fig. 3a), highlighting, as expected, the importance of this numerical component in YLD determination. The within environment correlation indicates the overall differences in YLD that were closely linked to genotypic variations in GN. On the other hand, the correlation between YLD and GW_1000_ was weak within and across environments (Fig. 3b). While there was a significant positive correlation between YLD and GW_1000_ when analyzing the entire dataset, the significance of this correlation differed when considering individual environments. The correlation was not significant in P13, slightly significant in MJ15, and highly significant in P15 (Fig. 3b). Regarding Sa, TBa, PS, Sm, TBm and HI the genotype generally explained a significant portion of the variation in the traits, while the environment and genotype x environment interactions varied in their contributions (Supplementary Table 2).

**Figure 3.**
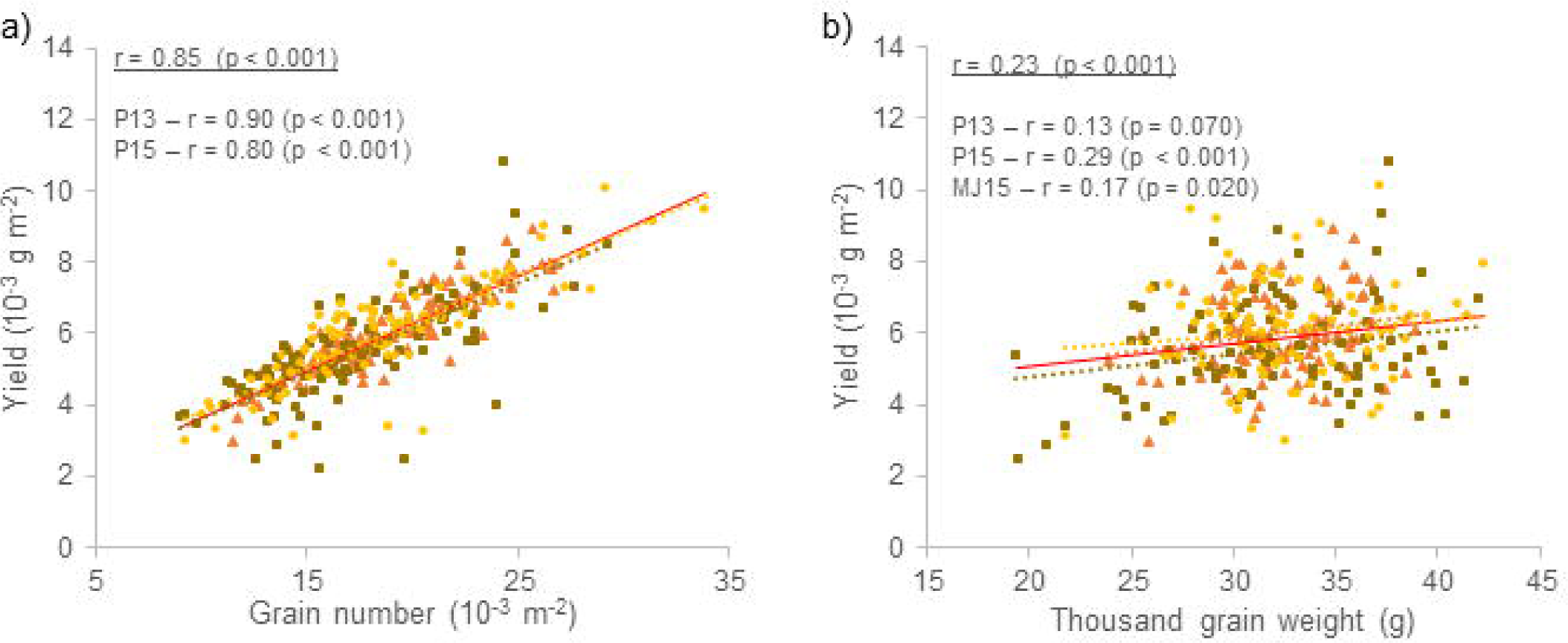
Relationship between grain yield vs grain number per m^-2^ and thousand grain weight between (solid red line) and within environments: P13 (orange triangles and dotted line), P15 (brown squares and dotted line) and MJ15 (yellow circles and dotted line). Pearson correlations and p-values for genotypes within each environment are indicated. The underlined r shows correlation across environments (E).

### 3.3. Relationships between grain number, thousand grain weight and the fruiting efficiencies

The variance in FEm and FE was significantly influenced by the environment, explaining 6% and 17% of the variance, respectively. Nevertheless, the genotype accounted for a larger proportion of the variance in FEm (39%) and FE (30%), indicating substantial genetic control over these traits. Notably, the genotype x environment interaction was not significant for either variable, although there was a discernible trend for FEm (Table 2).

**Table 2.** Percentage of total variation (considering the sum of squares) and p-values for the environment, genotype, and genotype x environment interaction.

| Trait | Environment | Genotype | Genotype X Environment |
| --- | --- | --- | --- |
| FEm | 6 | 39 | 24 |
|  | < 0.0001 | 0.0037 | 0.0722 |
| FE | 17 | 30 | 24 |
|  | < 0.0001 | < 0.0001 | 0.4040 |

FEm: fruiting efficiency at maturity (grains $g_{CH}^{-1}$ ), FE: fruiting efficiency (grains $g_{SDW}^{-1}$ )

Phenotypic associations between GN and FEm as well as FE were analyzed within and across environments. GN exhibited a strong positive correlation with both FEm and FE within and across environments (Fig. 4a-b), indicating that higher GN were associated with greater efficiencies in FEm and FE. In contrast, GW_1000_ showed a significant negative correlation with FEm and FE across environments (r = -0.22, p = 0.004 and r = -0.19, p < 0.001, respectively), suggesting that greater FEm and FE were associated with lowerGW_1000_. However, the correlations between GW_1000_ and FEm or FE within individual environments were not consistently significant, indicating that this relationship varied depending on the specific environmental conditions (Fig. 4c-d).

**Figure 4.**
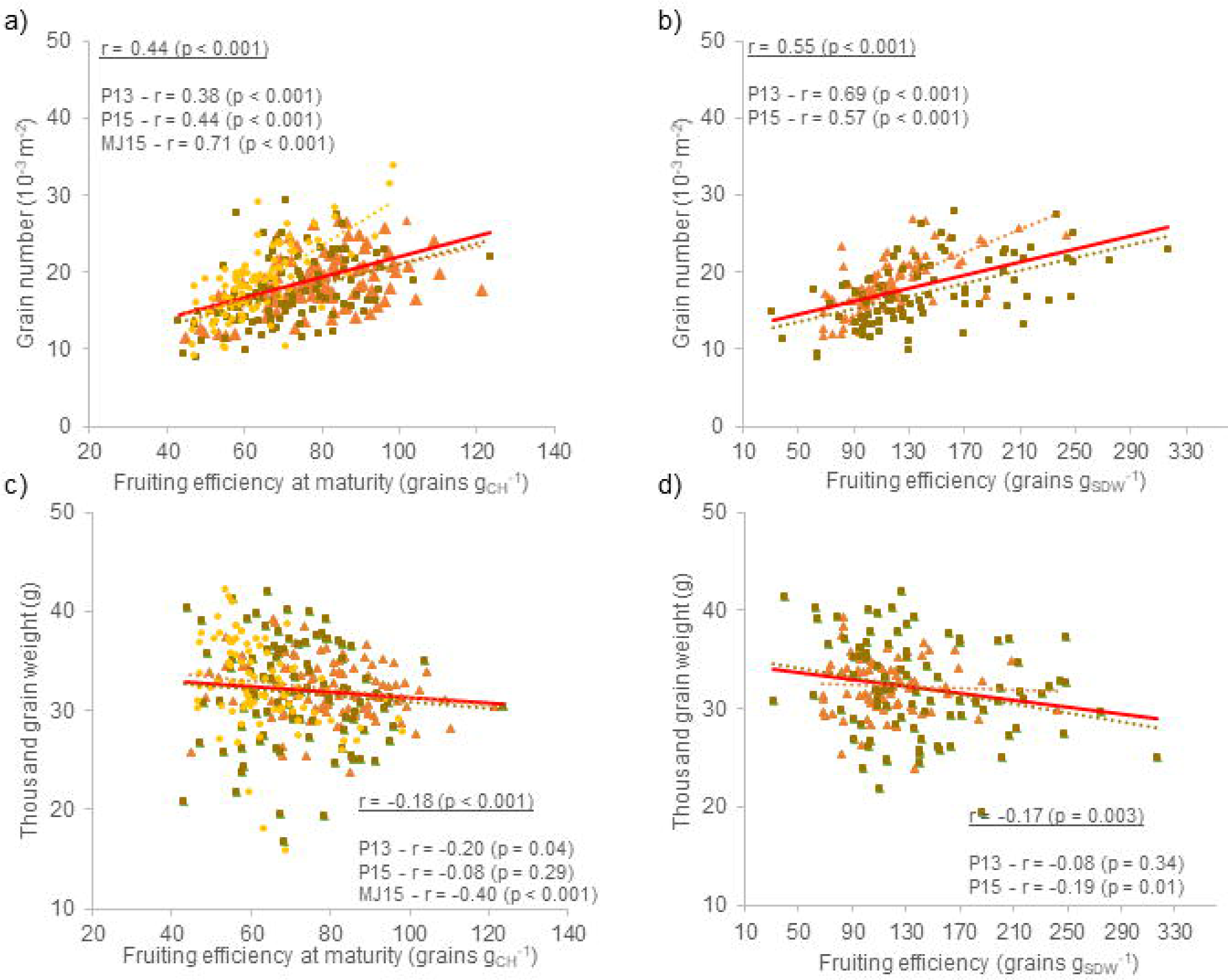
Relationship between grain number per m^-2^ and thousand grain weight vs fruiting efficiency and fruiting efficiency at maturity for two and three environments respectively: P13 (orange triangles and dotted line), P15 (brown squares and dotted line) and MJ15 (yellow circles and dotted line). Pearson correlations and p-values for genotypes within each environment are indicated. The underlined r shows correlation across environments (E).

### 3.4. QTL effect on yield and related traits

The effect of each QTL, namely *QFEm.perg-3A* and *QFEm.perg-5A*, as well as their two-way interaction, on various traits at the plot-level were carried out (Table 3 and Table 4). In the epistatic analysis between both QTL (*QFEm.perg-3A*\**QFEm.perg-5A*), significant epistatic interactions were observed only for SDW (p = 0.028) and GNs (p = 0.033) (Table 3, Fig. 5). For SDW, when the B19 allele was fixed in *QFFE.perg-5A*, the presence of the B2002 allele in the *QFEm.perg-3A* significantly increased SDW compared to the presence of B19 allele. However, when B2002 was fixed in *QFFE.perg-5A,* there was no significant differences in SDW between the B2002 and B19 allele in *QFEm.perg-3A* (Fig. 5a). For GNs, when the B19 allele was fixed in *QFFE.perg-5A*, there was no significant difference between the presence of the B2002 or B19 allele in *QFEm.perg-3A*. However, when the B2002 allele was fixed in *QFFE.perg-5A*, there was a significant increase in the GNs when the B19 allele was present in the *QFEm.perg-3A* compared to the presence of the B2002 allele (Fig. 5b). That is, when the B19 allele was present in either the *QFEm.perg-*3A or the *QFFE.perg-*5A, the GNs was approximately 45-47 grains per spike. However, when the B2002 allele was present in both QTL, GNs decreased to 42 grains per spike.

**Figure 5.**
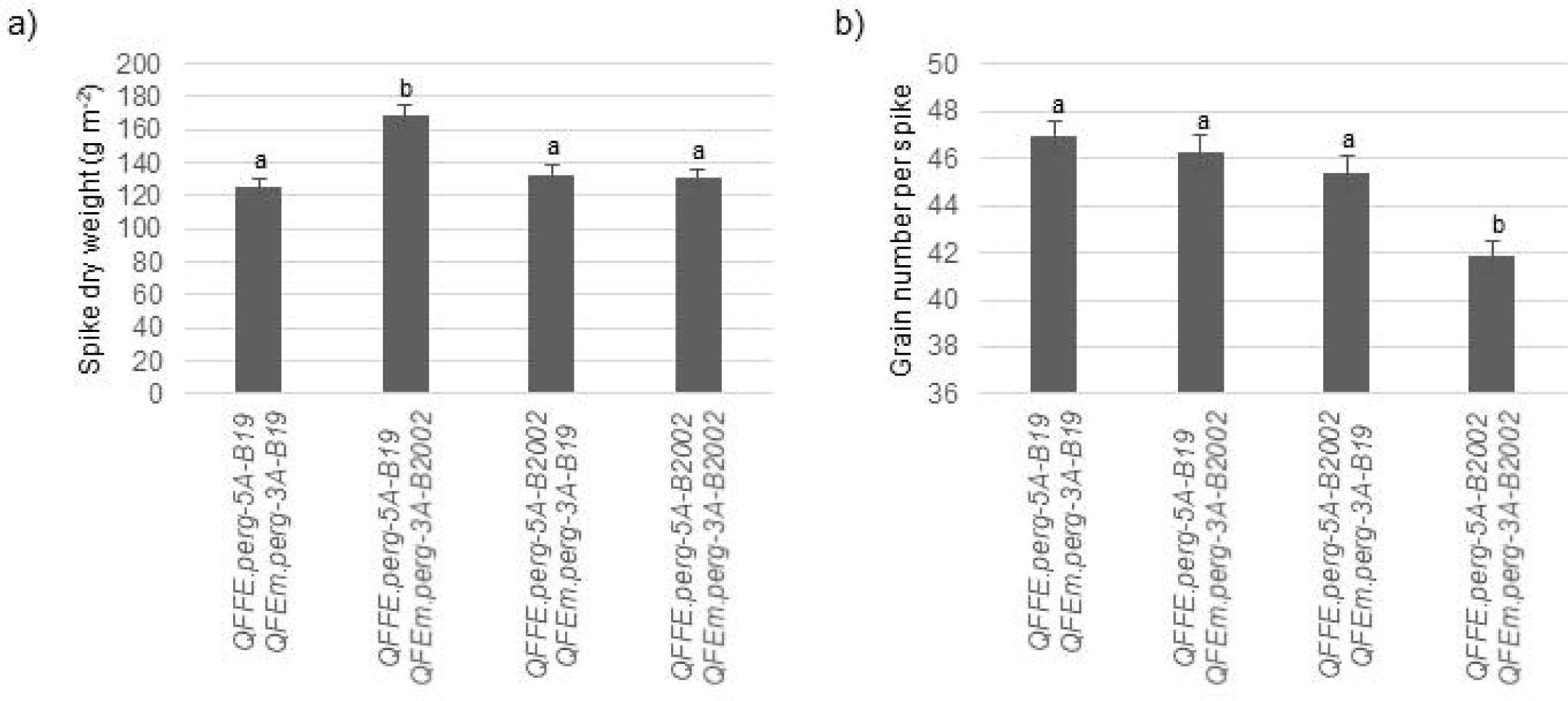
Two-way interaction plots between *QFFE.perg-5A* and *QFEm.perg-3A* on SDW and GNs. Different letters indicate significant differences (Tukey test α =0.05). The bars display standard errors.

**Table 3.**
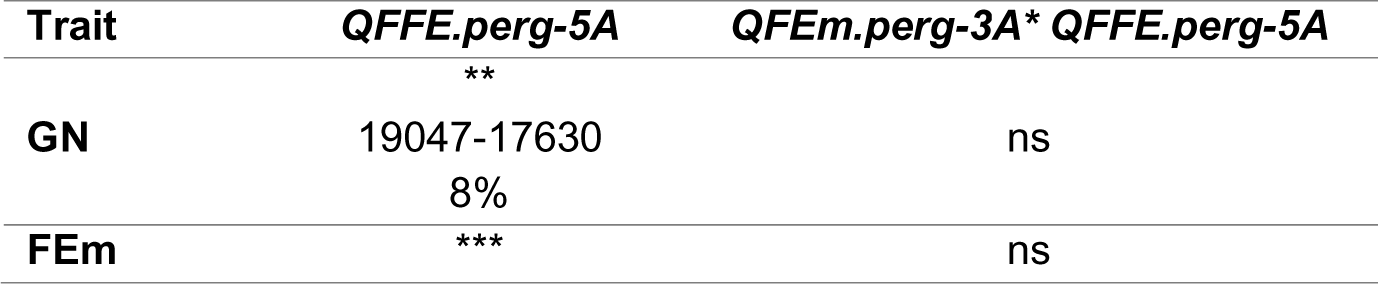
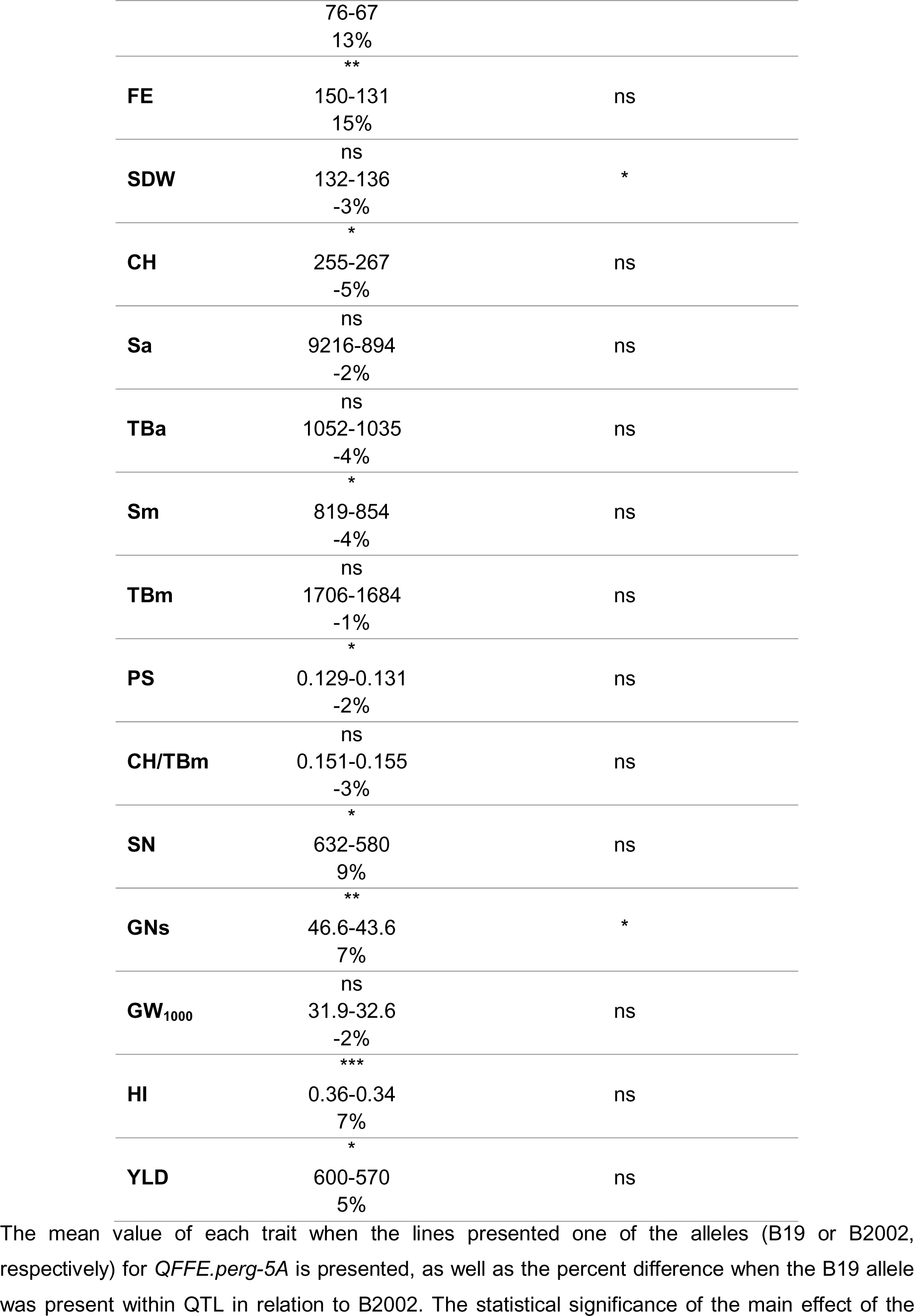
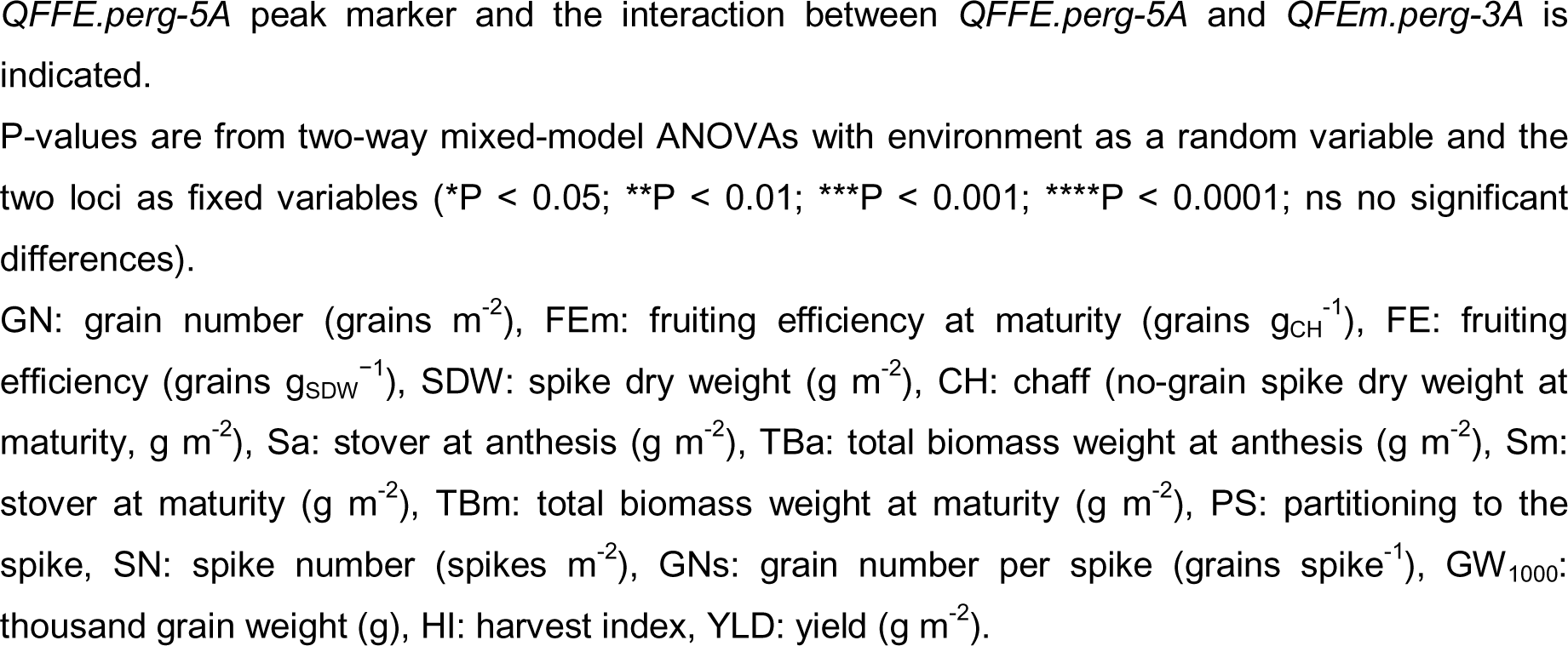
Individual effect of *QFFE.perg-5A* and *QFEm.perg-3A* \**QFEm.perg-5A* on plot-level traits.

**Table 4.** Mean values of yield and related traits for *QFEm.perg-3A* on each environment.

| Plot-level trait | Environment |  |  |  |
| --- | --- | --- | --- | --- |
|  | *QFEm.perg-3A* allele | P13 | P15 | MJ15 |
| GN | B19 | 18715 | 18110 | **20156** |
|  | B2002 | 18720 | 17029 | **16874** |
| FEm | B19 | 82.2 | 73.1 | **66.4** |
|  | B2002 | 77.6 | 71.2 | **59.0** |
| FE | B19 | 114.8 | 144.0 | - |
|  | B2002 | 111.0 | 135.2 | - |
| **SDW** | B19 | 163 | 129 | - |
|  | B2002 | 167 | 136 | - |
| **CH** | B19 | 233 | 255 | 298 |
|  | B2002 | 247 | 244 | 290 |
| **Sa** | B19 | 918 | 897 | - |
|  | B2002 | 943 | 900 | - |
| **TBa** | B19 | 1090 | 1030 | - |
|  | B2002 | 1101 | 1035 | - |
| **Sm** | B19 | 802 | 874 | 861 |
|  | B2002 | 839 | 817 | 813 |
| **TBm** | B19 | 1644 | 1693 | 1793 |
|  | B2002 | 1692 | 1635 | 1690 |
| **PS** | B19 | 0.15 | 0.13 | - |
|  | B2002 | 0.15 | 0.13 | - |
| **CH/TBm** | B19 | 0.14 | 0.15 | 0.17 |
|  | B2002 | 0.14 | 0.15 | 0.17 |
| **SN** | B19 | 628 | 525 | **635** |
|  | B2002 | 638 | 494 | **581** |
| **GNs** | B19 | **37.3** | 57.4 | **44.0** |
|  | B2002 | **35.6** | 55.2 | **41.0** |
| **GW<sub>1000</sub>** | B19 | **31.8** | 30.8 | **31.8** |
|  | B2002 | **32.8** | 32.1 | **34.0** |
| **HI** | B19 | 0.36 | 0.34 | 0.35 |
|  | B2002 | 0.36 | 0.34 | 0.34 |
| **YLD** | B19 | 589 | 542 | **631** |
|  | B2002 | 603 | 549 | **581** |

Black values indicate differences in the ANOVA ( $p < 0.05$ ). P13: Pergamino 2013, P15: Pergamino 2015, MJ15: Marco Juárez 2015.
GN: grain number (grains $m^{-2}$ ), FEm: fruiting efficiency at maturity (grains $g_{CH}^{-1}$ ), FE: fruiting efficiency (grains $g_{SDW}^{-1}$ ), SDW: spike dry weight ( $g\ m^{-2}$ ), CH: chaff (no-grain spike dry weight at maturity, $g\ m^{-2}$ ), Sa: stover at anthesis ( $g\ m^{-2}$ ), TBa: total biomass weight at anthesis ( $g\ m^{-2}$ ), Sm: stover at maturity ( $g\ m^{-2}$ ), TBm: total biomass weight at maturity ( $g\ m^{-2}$ ), PS: partitioning to the spike, SN: spike number (spikes $m^{-2}$ ), GNs: grain number per spike (grains spike $^{-1}$ ), GW<sub>1000</sub>: thousand grain weight (g), HI: harvest index, YLD: yield ( $g\ m^{-2}$ ).

It was observed that *QFFE.perg-5A* exhibited no interaction with the environment across all the traits under study (Supplementary Table 3). Therefore, the individual effect of *QFFE.perg-5A* in the whole dataset was analyzed (Table 3). When the B19 allele was present, GN increased by 8% due to higher FEm and FE (13% and 15%, respectively), resulting in 1417 more grains per m^2^ (Table 3) in spite of a slight but non-significant (p = 0.5241) trend towards a decrease in SDW by -3% (and this trait responded to the previously described epistatic interaction). In contrast, the CH showed a significant difference between both alleles (p = 0.0298) (Table 3), reducing by 5% when the B19 allele was present. The stover at anthesis (Sa) and the total biomass at anthesis (TBa) were not affected by the presence of the B19 allele (Table 3). While the stover at maturity (Sm) was reduced, the total biomass at maturity (TBm) was not affected. Despite a slight reduction in partitioning of dry matter to the spikes at anthesis (PS), the relationship between CH and TBm was not modified (Table 3). These results were associated with an increase of 52 spikes per m^-2^ (9%) when the B19 allele was present. Consequently, the greater GN was achieved not only by the increase in GNs by 7% but also in the SN (Table 3). The GW_1000_ was slightly reduced by the presence of this allele (2%), though it was not significantly (p = 0.1111). This resulted in a significant 7% increase in the harvest index (HI) and a 5% increase in yield (Table 3).

The *QFEm.perg-3A* showed a significant interaction with the environment for GN, SN and Sm (Supplementary Table 3). Given the importance of these traits on YLD, the effect of *QFEm.perg-3A* on each environment was separately analyzed (Table 4). In P13, a slight increase in GNs (p = 0.0418) associated with a reduction in GW_1000_ (p = 0.0278) was observed, but no difference in YLD, FEm, or FE was found in the presence of any allele. In P15, no significant difference was found in any of the traits under study (Table 4). On the other hand, in MJ15 several traits showed significant differences depending on the presence of the different alleles. The SN and GNs were notably higher when the B19 allele was present, resulting in greater GN (3282 more grains per m^-2^) (Table 4). This was consequence of a highly significant difference in FEm (11%) which resulted in a significant response in YLD (8%), despite a significant decrease in GW_1000_ (7%) when B19 was present (Table 4).

## Discussion

The integration of marker-assisted selection (MAS) into traditional breeding methods has significantly improved breeding accuracy. With the increasing availability of molecular markers and genetic maps, MAS has become a viable strategy for targeting QTL. Specifically, the use of molecular markers identified at the spike level and their extrapolation at the plot level provides a robust approach for unraveling the genetic control underlaying spike-related traits. Understanding the ecophysiological aspects of wheat yield, considering the possible trade-off between some of the numerical components is crucial for interpreting our results. The integration of this aspects not only enhances our understanding of the genetic control but also accelerates breeding programs and ultimately improve crop productivity.

To the best of our knowledge, there has been no prior investigation into the plot level impact of QTL identified at the spike level in wheat, at least for QTL for GNs, FEms or FEs. In previous studies, our team conducted QTL analysis at the spike level to identify genetic loci associated with spike fertility and related traits (Pretini et al., 2021b). Furthermore, we successfully validated these QTL associations with spike fertility in a separate study (Pretini et al., 2020b). In the current work, our objective was to examine the impact of these QTL at the plot level. Drawing from spike-level data, as elucidated by Pretini et al. (2021a), it becomes apparent that three distinct spike ideotypes can be characterized, each differing in fruiting efficiency (FE). The first ideotype (ID1) is associated to low FE, while ID2 and ID3 exhibit high FE. Differences between ID1 and ID2 primarily correspond to the variations in fertile florets per spikelet, with no additional variations in the spike structure or SDW (Guo et al., 2017; Sakuma et al., 2019). On the other hand, differences between ID1 and ID3, manifest as greater number of fertile florets and grains per spikelet, accompanied by a reduction in spike structure and SDW (Alonso et al., 2018; Pretini et al., 2020a, b).

In our current investigation, we observed that the presence of the B19 allele in *QFFE.perg-5A* led to an increase in GN, consistent with the findings at the individual spike-level where greater GNs was observed (Pretini et al., 2020b). This improvement in GN was associated with an increase in FE and FEm. The SDW showed a trend to reduce (when the B19 allele was present also in the *QFEm.perg-3A*), while the CH slightly decreased by the presence of this allele, contrary to the observations obtained at spike-level where reductions in SDWs (-6%) and CHs (-5%) were significant (Pretini et al., 2020b) and no epistasis was observed between QTL. This discrepancy in results between the plot-level and spike-level analyses could be explained by the increase of SN, which may compensate for the reduction of the individual spike dry weight and help to achieve similar spike dry weight per unit area (SDW). As the dry matter partitioning to reproductive structures at anthesis (PS) was slightly affected by the allele’s presence, a greater number of lighter spikes were set. Whenever the allele from B19 was present in *QFFE.perg-5A* the GNs was higher, highlighting the importance of this allele in determining this trait. Although the presence of the B19 allele in one or both QTLs did not show statistically significant differences, it tended to present more GNs when it was present in both QTL. Additionally, having the allele present in only one QTL was enough to result in a statistical difference in GNs. As a result, the greater GN achieved when the B19 allele was present at *QFFE.perg-5A* was consequence not only of the increase in GNs but also on the SN, reinforcing the importance of the plot level studies when assessing the genetic control of yield-related traits. The GW_1000_ was not modified by the presence of this allele in *QFFE.perg-5A*, showing only a trend to lighter grains, which contrast the observations obtained at individual spike-level, where the grain weight was significantly reduced (Pretini et al., 2020b). Overall this resulted in a yield increment slightly higher than that at the individual spike level (Pretini et al., 2020b), which was associate to greater harvest index (HI) with no difference in the total crop growth (TBm).

Our plot-level results revealed a significant interaction between *QFEm.perg-3A* and the environment, particularly influencing GN, SN, and Sm. The analysis of *QFEm.perg-3A*’s impact on each environment revealed marginal differences. In P15, there were no significant differences in any of the traits under study, whereas in P13, differences were observed only in GW_1000_ and GNs, with no impact on YLD. On the other hand, the presence of the B19 allele in MJ15 showed significant differences in YLD and several related traits. Additionally, Pretini et al. (2020b) observed, at the spike-level, a reduced impact of *QFEm.perg-3A* during anthesis, with a more pronounced association with grain set (GST) than with fertile florets per spikes (FF). The lack of association of the *QFEm.perg-3A* to the determination of fertile florets, which is the main determinant of grain number in wheat (Siddique et al., 1989; González et al., 2003, 2005) may explain the variable impact of the QTL.

The positive impact of *QFFE.perg-5A* at the plot-level reinforced the findings reported by Pretini et al. (2020b) at the spike-level. The allele identified by Pretini et al. (2020b) positively impacted FE and increased not only the grains at spike-level (GNs) but also those per unit area (GN). Importantly, this enhancement was not associated with a decrease in SN, a situation that might have occurred if only GNs had been considered. In the Gaju et al. (2009) study, for example, a reduction in SN was observed in the one of the large-spike phenotype lines carrying higher GNs, which resulted in reduced GN and YLD. On the contrary, our investigation revealed that the presence of the favorable alleles enhancing FE led to a higher SN, without modifying the SDW. This observation could be explained by the spike ideotype concept we discussed previously, where the spikes characterized by a high coefficient entail lesser dry matter investment in spike structure (ID3). Consequently, under the same SDW partition (PS), these ideotypes could sustain or even augment the SN.

Understanding the effect and location of a marker is crucial for yield improvement; however, its integration into a breeding program may not be relevant if the marker lacks variability in the utilized germplasm or has already been fixed in the new released cultivars. To address this concern, we analyzed a total of 123 commercial varieties adapted to the Rolling Pampas in Argentina and in Uruguay, released between 2000 and 2024. These varieties comprise germplasm from the International Maize and Wheat Improvement Center (CIMMYT) as well as those with French origins, developed by 14 different companies^1^. Interestingly, our analysis revealed that the favorable allele (B19) of the *QFFE.perg-5A* was present in only 24% of the studied cultivars, suggesting its incorporation could be a potential tool for increasing yield.

Ongoing progress in developing of high-density mapping population will enable the identification of candidate genes, thereby enhancing our understanding of the fine molecular and physiological processes involved.

## Abbreviations

B2002: BioINTA2002
B19: Baguette 19
CH: chaff (no-grain spike dry weight at maturity, g m^-2^)
DH: double haploid
FE: fruiting efficiency (grains g_SDW_^−1^)
FEm: fruiting efficiency at maturity (grains g_CH_^-1^)
FEs: fruiting efficiency at spike level (grains g_SDW_^−1^)
FEms: fruiting efficiency at maturity at spike level (grains g ^-1^)
FFs: fertile florets per spike (number spike^-1^)
FFE: fertile floret efficiency (florets g_SDW_^-1^)
GN: grain number (grains m^-2^)
GNs: grain number per spike (grains spike^-1^)
GW: grain weight per spike (g spike^-1^)
GW_1000_: thousand grain weight (g)
HI: harvest index
PS: partitioning to the spike
QTL: quantitative trait loci
RCBD: randomized complete block design
SDW: spike dry weight (g m^-2^)
SDWs: spike dry weight per spike (g spike^-1^)
Sa: stover at anthesis (plant weight without spike weight, g m^-2^)
Sm: stover at maturity (plant weight without spike and grain weights, g m^-2^)
SN: spike number (spikes m^-2^)
TBa: total biomass weight at anthesis (g m^-2^)
TBm: total biomass weight at maturity (g m^-2^)
YLD: yield (g m^-2^)

## Author contribution statement

IIT and FGG carried out the phenotyping experiments during 2012 and 2013. NP, IIT and FGG carried out the phenotyping experiments during 2015. NP realized the QTL detection and wrote the first manuscript with revision from LSV, PS and FGG. NP, GF and PS characterized wheat lines for presence/absence of marker alleles. FGG designed and coordinated the project.

## Acknowledgments

The present work was funded by the Agencia Nacional de Promoción de la Investigación, el desarrollo tecnológico y la innovación (Agencia), Argentina (PICT 2012-1198, PICT 2014-1283, PICT 2019-3256, PICT 2021-IIIA-0122), the Instituto Nacional de Tecnología Agropecuaria (INTA, PD-I102, PEI-109) Argentina, the Monsanto Beachell-Bourlag Scholarship, the Universidad nacional del Noroeste de la Provincia de Buenos Aires (UNNOBA, SIB 2015, SIB 2017, SIB 2019) Argentina and the EU FP7 Funding (ADAPATWHEAT 289842). GF is research fellow the Agencia.

## Footnotes

1 (Nidera Seeds, Argentina; Asociacion de Cooperativas Argentinas, SAS Florimond Deprez Veuve et Fils, Asociados Don Mario SA, Or Melhoramiento de Sementes Ltda, Instituto Nacional de Tecnología Agropecuaria, Buck Semillas SA, Biotrigo Genetida Ltda, Instituto Paraguayo de Tecnología Agropecuaria, Criadero Klein SA, Limagrain, Bioseminis, Syngenta Crop Protection Ag, BO Paiva)

## References

Abbate PE, Andrade FH, Lázaro L, Bariffi JH, Berardocco HG, Inza VH, Marturano F. 1998. Grain yield increase in recent Argentine wheat cultivars. Crop Science 38, 1203–1209.

Alonso MP, Panelo J, Mirabella NE, Pontaroli AC. 2018. Selection for high spike fertility index increases genetic progress in grain yield and stability in bread wheat. Euphytica 214:112.

Basile SML, Ramirez IA, Crescente JM, Conde MB, Demichelis M, Abbate PE, Rogers WJ, Pontaroli AC, Helguera M, Vanzetti LS. 2019. Haplotype block analysis of an Argentinean hexaploidy wheat collection and GWAS for yield components and adaptation. BMC Plant Biology 19:553.

Borlaug NE. 2007. Sixty-two years of fighting hunger: personal recollections. Euphytica 157(3):287–97.

Chand R. 2009. Challenges to ensuring food security through wheat. CAB reviews: Perspectives in agriculture, veterinary science, nutrition and natural resources. 4(065):1–13.

Chen H, Iqbal M, Yang RC, Spaner D. 2016. Effect of Lr34/Yr18 on agronomic and quality traits in a spring wheat mapping population and implications for breeding. Molecular Breeding 36:5.

Chen H, Semagn K, Iqbal M, Moakhar NP, Haile T, N’Diaye A, Yang R-C, Hucl P, Pozniak C, Spaner D. 2017. Genome-wide association mapping of genomic regions associated with phenotypic traits in Canadian western spring wheat. Molecular Breeding 37:141.

Cheng R, Doerge RW, Borevitz J. 2017. Novel resampling improves statistical power for multiple-trait QTL mapping. G3-Genes Genomes Genetics 7:813.

Cobb JN, Biswas PS, Platten JD. 2019. Back to the future: Revisiting MAS as a tool for modern plant breeding. Theoretical and Applied Genetics 132, 647–667.

Deng Z, Cui Y, Han Q, Fang W, Li J, Tian J. 2017. Discovery of consistent QTLs of wheat spike-related traits under nitrogen treatment at different development stages. Frontiers Plant Science 8:2120.

Di Rienzo JA, Casanoves F, Balzarini MG, González L, Tablada M, Robledo CW. 2016. Grupo InfoStat, FCA, Universidad Nacional de Córdoba, Argentina.

Dreccer M, van Herwaarden A, Chapman S. 2009. Grain number and grain weight in wheat lines contrasting for stem water soluble carbohydrate concentration. Field Crops Research 112, 43–54.

Dreccer MF, Chapman SC, Rattey AR, Neal J, Song Y, Christopher JJ, Reynolds M. 2013. Developmental and growth controls of tillering and water-soluble carbohydrate accumulation in contrasting wheat (Triticum aestivum L.) genotypes: can we dissect them? Journal of Experimental Botany 64(1), 143–60.

FAO, IFAD, UNICEF, WFP and WHO. 2021. The State of Food Security and Nutrition in the World 2021. Transforming food systems for food security, improved nutrition and affordable healthy diets for all. Rome, FAO.

Fischer RA. 1975. Yield potential in a dwarf spring wheat and the effect of shading 1. Crop Science 15:607–613.

Fischer RA. 1983. Wheat. Potential productivity of field crops under different environments. International Rice Research Institute, Los Baños pp 129–154.

Fischer RA. 1985. Number of kernels in wheat crops and the influence of solar radiation and temperature. Journal of Agricultural Science 105:447–461.

Fischer RA, Connor DJ. 2018. Issues for cropping and agricultural science in the next 20 years. Field Crops Research 222, 121–142.

Fischer RA, Rebetzke GJ. 2018. Indirect selection for potential yield in early-generation, spaced plantings of wheat and other small-grain cereals: a review. Crop Pasture Science 69(5):439.

Gaju O, Reynolds MP, Sparkes DL, Foulkes MJ. 2009. Relationships between large-spike phenotype, grain number, and yield potential in spring wheat. Crop Science 49, 961–973.

Gerard GS, Alqudah A, Lohwasser U, Börner A, Simón MR. 2019. Uncovering the genetic architecture of fruiting efficiency in bread wheat: a viable alternative to increase yield potential. Crop Science 59:1–17.

Ghiglione HO, Gonzalez FG, Serrago R, Maldonado SB, Chilcott C, Cura JA, Miralles DJ, Zhu T, Casal JJ. 2008. Autophagy regulated by daylength sets the number of fertile florets in wheat. Plant Journal 55:1010–1024.

González FG, Slafer GA, Miralles DJ. 2003. Grain and floret number in response to photoperiod during stem elongation in fully and slightly vernalized wheats. Field Crops Research 81, 17–27

González FG, Slafer GA, Miralles DJ. 2005. Photoperiod during stem elongation in wheat: is its impact on fertile floret and grain number determination similar to that of radiation? Functional Plant Biology 32, 181–188.

González FG, Miralles DJ, Slafer GA. 2011a. Wheat floret survival as related to pre-anthesis spike growth. Journal of Experimental Botany 62:4889–4901.

González F, Terrile II, Falcón MO. 2011b. Spike fertility and duration of stem elongation as promising traits to improve potential grain number (and yield): variation in modern Argentinean wheats. Crop Science 51:1693.

Guo Z, Chen D, Alqudah AM, Röder MS, Ganal MW, Schnurbusch T. 2017. Genome-wide association analyses of 54 traits identified multiple loci for the determination of floret fertility in wheat. New Phytologyst 214:257–270.

Isham K, Wang R, Zhao W, Wheeler J, Klassen N, Akhunov E, Chen J. 2021. QTL mapping for grain yield and three yield components in a population derived from two high-yielding spring wheat cultivars. Theoretical and Applied Genetics 134(7):2079–2095.

Kirby EJM. 1988. Analysis of leaf, stem and ear growth in wheat from terminal spikelet stage to anthesis. Field Crop Research 18:127–140.

Mahibbur RM, Govindarajulu Z. 1997. A modification of the test of Shapiro and Wilk for normality. Journal of Applied Statistics 24(2):219–35.

Mammadov J, Aggarwal R, Buyyarapu R, Kumpatla. 2012. SNP markers and their impact on plant breeding. International Journal of Plant Genomics, 728398.

Martino DL, Abbate PE, Cendoya MG, Gutheim F, Mirabella NE, Pontaroli AC, Saranga Y 2015. Wheat spike fertility: inheritance and relationship with spike yield components in early generations. Plant Breeding 134(3):264–270.

Sakuma S, Golan G, Guo Z, et al. 2019. Unleashing floret fertility in wheat through the mutation of a homeobox gene. Proceedings of the National Academy of Sciences, USA 116, 5182.

Pang Y, Liu C, Wang D, et al. 2020. High-Resolution Genome-wide Association Study Identifies Genomic Regions and Candidate Genes for Important Agronomic Traits in Wheat. Molecular Plant 13, 1311–27.

Pretini N, Terrile II, Gazaba LN, Donaire G, Arisnabarreta S, Vanzetti LS, González FG. 2020a. A comprehensive study of spike fruiting efficiency in wheat. Crop Science 1–15.

Pretini N, Vanzetti LS, Terrile II, Börner A, Plieske J, Ganal M, Röder M, González FG. 2020b. Identification and validation of QTL for spike fertile floret and fruiting efficiencies in hexaploid wheat (*Triticum aestivum* L.). Theoretical and Applied Genetics 133, 2655–2671.

Pretini N, Alonso MP, Vanzetti LS, Pontaroli AC, González FG. 2021a. The physiology and genetics behind fruiting efficiency: a promising spike trait to improve wheat yield potential. Journal of Experimental Botany 72, 11:3987–4004.

Pretini N, Vanzetti LS, Terrile II, Donaire G, González FG. 2021b. Mapping QTL for spike fertility and related traits in two doubled haploid wheat (*Triticum aestivum L.*) populations. BMC Plant Biology 21, 353.

Quraishi UM, Murat F, Abrouk M, Pont C, Confolent C, Oury FX, Ward J, Boros D, Gebruers K, Delcour JA, Courtin CM, Bedo Z, Saulnier L, Guillon F, Balzergue S, Shewry PR, Feuillet C, Charmet G, Salse J. 2011. Combined meta-genomics analyses unravel candidate genes for the grain dietary fiber content in bread wheat (*Triticum aestivum L*.). Functional & Integrative Genome 11:71–83.

Reynolds M, Foulkes J, Furbank R, Griffiths S, King J, Murchie E, Parry M, Slafer G. 2012. Achieving yield gains in wheat. Plant Cell Environment 35:1799–1823.

Siddique KHM, Kirby EJM, Perry MW. 1989. Ear:stem ratio in old and modern wheat varieties; relationship with improvement in number of grains per ear and yield. Field Crops Research 21, 59–78.

Stapper M, Fischer RA. 1990. Genotype, sowing date and plant spacing influence on high-yielding irrigated wheat in southern New South Wales. II. Growth, yield and nitrogen use. Australian Journal Agricultural Research 41(6):1021–1041.

Terrile II, Miralles DJ, González FG. 2017. Fruiting efficiency in wheat (*Triticum aestivum L*): Trait response to different growing conditions and its relation to spike dry weight at anthesis and grain weight at harvest. Field Crops Research, 201, 86–96.

Xu YF, Li SS, Li LH, Ma FF, Fu XY, Shi ZL, Xu HX, Ma PT, An DG. 2017. QTL mapping for yield and photosynthetic related traits under different water regimes in wheat. Molecular Breeding 37:34.

Yu M, Mao SL, Hou DB, Chen GY, Pu ZE, Li W, Lan XJ, Jiang QT, Liu YX, Deng M, Wei YM. 2018. Analysis of contributors to grain yield in wheat at the individual quantitative trait locus level. Plant Breeding 137:35–49.

Zadoks JC, Chang TT, Konzak CF. 1974. A decimal code for the growth stages of cereals. Weed Research 14:415–42.

Zhai H, Feng Z, Du X et al. 2018. A novel allele of TaGW2-A1 is located in a finely mapped QTL that increases grain weight but decreases grain number in wheat (*Triticum aestivum L*.). Theoretical and Applied Genetics 131:539–553.

Zheng T, Hua C, Li L, Sun Z, Yuan M, Bai G, Humphreys G, Li T. 2021. Integration of meta-QTL discovery with omics: Towards a molecular breeding platform for improving wheat resistance to Fusarium head blight. Crop Journal 9, 739–749.

